# A conserved PLK1 docking site in TopBP1 maintains genome integrity during mitosis

**DOI:** 10.1101/2022.03.30.486397

**Authors:** Jiayi Li, Jonas Bagge, Michael Lisby, Jakob Nilsson, Vibe H. Oestergaard

## Abstract

TopBP1 is a large scaffold protein with multiple functions in genome integrity. We previously identified a novel role for TopBP1 during M phase by showing that TopBP1 reduces carry-over of DNA damage to daughter cells. This function emerges as a critical backup pathway in BRCA deficient cells, yet many aspects of TopBP1 regulation during mitosis are unclear. The mitotic kinase PLK1 has been reported to interact with TopBP1 but the functional relevance of this is unclear. Here, we identify and characterize a conserved PLK1 docking site in TopBP1. Endogenous deletion of the PLK1 docking site in TopBP1 results in increased number of mitotic TopBP1 foci, increased DNA damage in daughter cells, deficient mitotic DNA repair synthesis and increased frequency of binucleation. At the same time, cell cycle distribution and ATR activation are normal in cells with the PLK1 docking site deletion in TopBP1. Interestingly, mutation of this site in TopBP1 renders cells sensitive to PARP inhibitors but not to camptothecin hinting to different cellular effects of the two chemotherapeutics. Altogether, our data indicate that the PLK1-TopBP1 interaction is critical for the mitotic function of TopBP1.

## Introduction

Maintaining genome integrity is important to avoid cell death and development of genetic diseases including cancers. Thus, multiple DNA repair pathways have evolved to protect the genome. DNA repair pathway choice is influenced by the type of DNA damage but also the cell cycle stage. In the recent years it has become clear that certain genome maintenance pathways are active in M phase. Moreover, cancer genomic studies also indicate that the mitotic repair pathways are error prone giving rise to structural variations (Dereli-Oz et al., 2011; Li et al., 2020). The mitotic genome maintenance pathways include mitotic DNA synthesis at underreplicated regions (MiDAS)(Bhowmick et al., 2016; Minocherhomji et al., 2015; Pedersen et al., 2015), processing of recombination intermediates (Balbo Pogliano et al., 2022; Matos and West, 2014), and tethering of DNA double-strand breaks to avoid missegregation of acentric chromosome fragments (Leimbacher et al., 2019). Interestingly, TopBP1 is key for all these processes (Balbo Pogliano et al., 2022; Leimbacher et al., 2019; Pedersen et al., 2015). TopBP1 is a large scaffold protein with no enzymatic function. It contains 9 BRCT domains named BRCT0 to BRCT8. It has essential roles both as an ATR activator via its ATR activation domain and as a key player in replication initiation (Bagge et al., 2020).

A major unresolved question is how TopBP1 functions are regulated during mitosis. TopBP1 is phosphorylated on multiple sites during mitosis and has also been found to interact with the mitotic kinase PLK1. However, whether the interaction with PLK1 is direct and of functional importance is presently unclear as no separation-of-function mutations in TopBP1 have been established.

PLK1 is a kinase that is highly active in and around mitosis. It is required for dampening checkpoint arrest at G2 to promote mitotic entry (van Vugt and Medema, 2005) and during mitosis it drives many mitotic processes including spindle assembly, regulation of the anaphase-promoting complex (APC), mitotic exit and cytokinesis (Joukov and De Nicolo, 2018).

PLK1 was shown to promote loading of RAD51 in interphase cells (Moudry et al., 2016) in a process that relies on BRCT7 and BRCT8 of TopBP1 for coordinating the PLK1-mediated phosphorylation of RAD51 (Moudry et al., 2016). Interestingly, PLK1-mediated RAD51-phoshorylation promotes MiDAS (Wassing et al., 2021). Furthermore, PLK1 activity is important for suppressing recruitment of NHEJ factors to camptothecin (CPT)-induced replication problems (Nakamura et al., 2021).

The polo-box domain of PLK1 is directing substrate recognition by binding to Serine-Threonine-Proline (STP)-like motifs, where the threonine has been phosphorylated by CDK or another proline-directed kinase (Reinhardt and Yaffe, 2013).

To understand how TopBP1 is regulated during mitosis we investigated the importance of a highly conserved STP site in TopBP1. Our results indicate that the STP site is a functional PLK1 docking site that mediates TopBP1-PLK1 interaction and is important for maintaining genomic integrity during M phase.

## Results

### PLK1 binds a conserved STP motif in TopBP1

PLK1 is known to bind to STP sequences that constitutes PLK1 docking motifs. Since PLK1 has been reported to bind TopBP1, we scanned vertebrate TopBP1 for such sequences. Indeed, we identified three STP sequences with the most N-terminal motif showing the highest degree of conservation (Fig. 1A). This STP motif is located close to BRCT2 in an unstructured region (Fig. 1A)(Day et al., 2018). To investigate if this motif can indeed interact with PLK1, we used *in vitro* isothermal titration calorimetry (ITC), which confirms a dissociation constant in the nanomolar to low micromolar range for the chicken and human TopBP1 phosphopeptides, respectively (Fig. 1B and supplementary figure 1A). Interestingly the PLK1 docking site in chicken TopBP1 is a double consensus site and our ITC measurements suggests that the more C-terminal motif binds with higher affinity (Fig. 1B).

**Figure 1.**
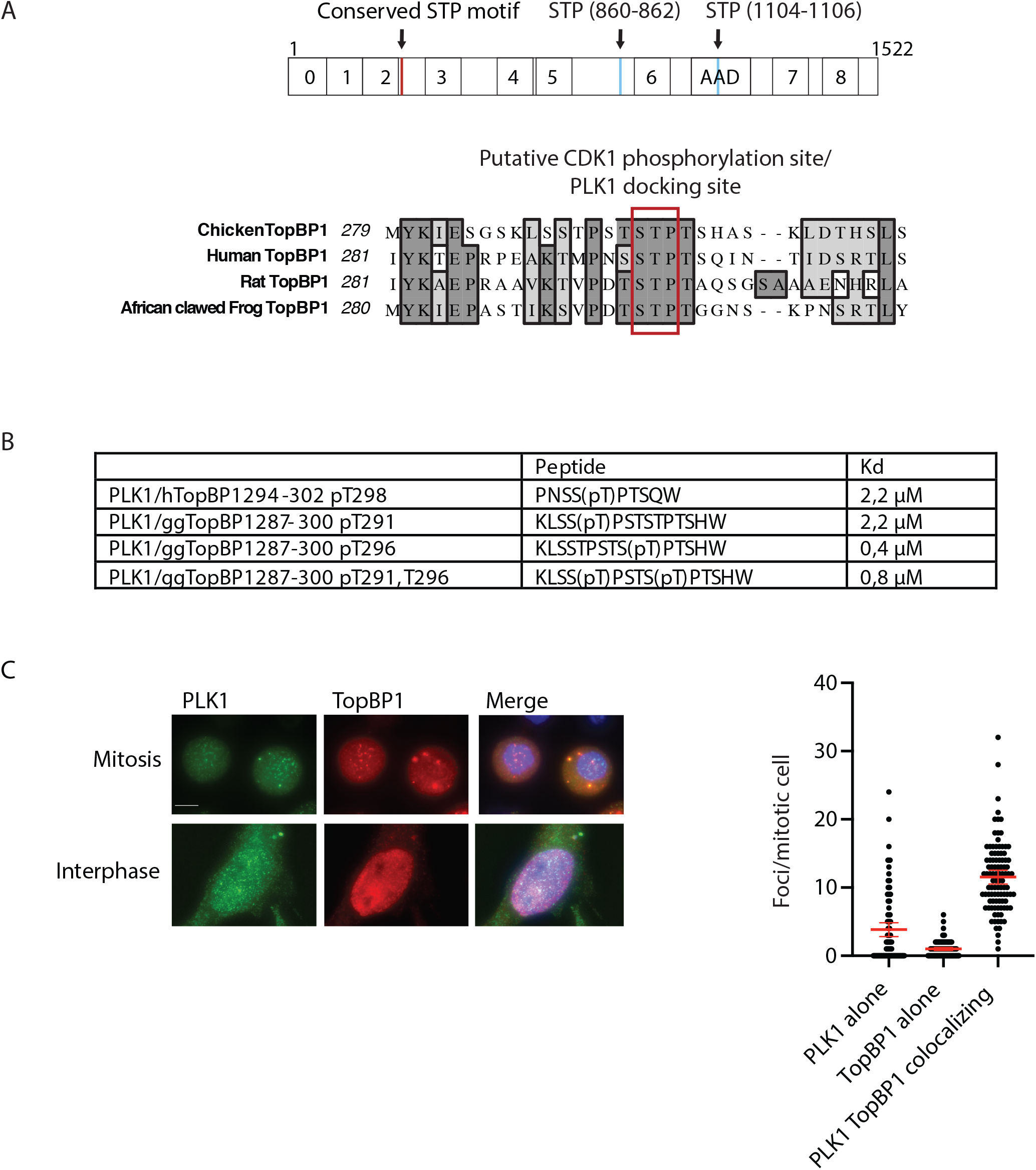
Characterization of a conserved PLK1 docking site in TopBP1. A. *Upper*, schematic representation of human TopBP1. BRCT domains are indicated with numbers. ATR activation domain is indicated (AAD). Red bar indicates the position of a conserved N-terminal STP. Blue bars indicate the positions of other conserved STP motifs. *Lower*, alignment of a putative PLK1 docking site in TopBP1 from the indicated vertebrate species. The STP sequence/putative PLK1 docking site is marked by a red frame. The dark grey shading indicates high conservation between different species. Light grey shadints indicate medium conservation of amino acids. B. Isothermal titration calorimetry data. Affinities and thermodynamic values of PLK1-TopBP1 peptide binding events inferred from ITC measurements performed at 25°C. Gibbs free energy (ΔG), enthalpy (ΔH), entropy (-TΔS), equilibrium dissociation constant (KD) and reaction stoichiometry (n) are shown. The affinity is defined by the Gibbs energy for binding ΔG = −RT lnKA = RT lnKD. The errors represent the standard error of the fitting. C. *Left*, Immunostaining of PLK1 and TopBP1 in HeLa cells during mitosis and interphase as indicated. Experiments were performed in duplicate and representative images are shown. *Right*, quantification of PLK1 and TopBP1 colocalizing foci as well as non-colocalization foci in immuno-stained mitotic HeLa cells. Error bars represent the means ± 95% CI.

TopBP1 interaction with PLK1 may give rise to cellular colocalization that can be studied by microscopy. To investigate whether TopBP1 colocalizes with PLK1, co-immunostaining of mitotic and interphase HeLa cells for PLK1 and TopBP1 was performed (Fig. 1C).

Quantitative analysis of the images reveals that TopBP1 foci and PLK1 foci overlap to a large extent in M phase (Fig. 1C). Notably, nearly all TopBP1 foci colocalizes with PLK1, whereas PLK1 seems to also form foci devoid of TopBP1 likely reflecting PLK1 localization to kinetochores. There is no obvious PLK1 and TopBP1 colocalization during interphase, indicating that the PLK1-TopBP1 complex might serve an important function in M phase.

### Deletion of the PLK1 docking site in TopBP1 does not affect ATR activation or cell cycle progression

To investigate the potential function of the PLK1 docking site that we identified, we set out to delete this site in endogenous TopBP1 in chicken DT40 cells, which we previously successfully used to decipher mitotic TopBP1 functions (Germann et al., 2014; Pedersen et al., 2015). We employed a CRISPR-Cas9 two-target-site strategy to remove a 54 base pair region encoding the STP motif resulting in an 18-amino-acid in-frame deletion (Fig. 2A).

**Figure 2.**
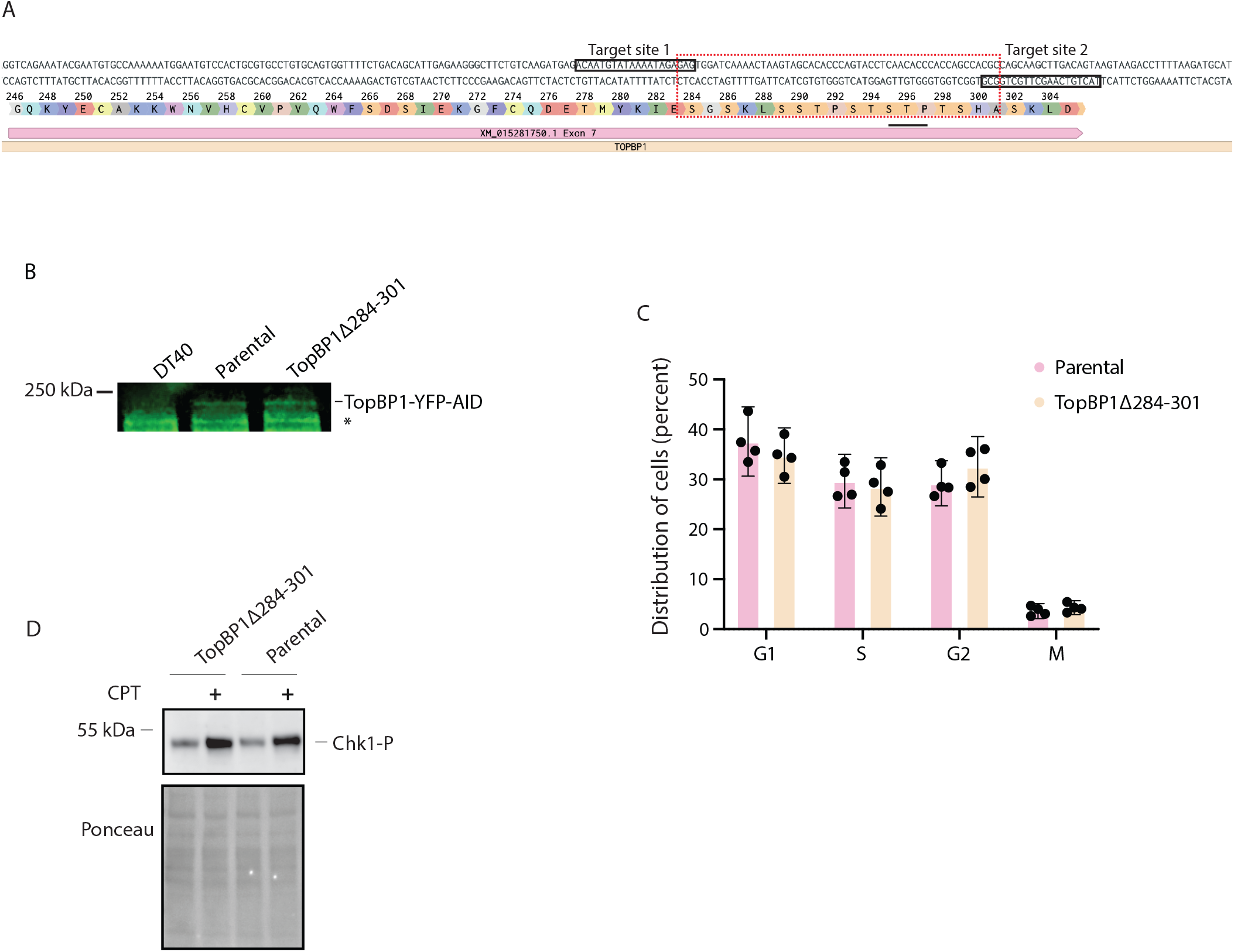
Endogenous deletion of the PLK1 docking in TopBP1. A. CRISPR targeting strategy for the endogenous deletion of a PLK1 docking site to generate TopBP1Δ284-301. B. Extracts from the indicated cell lines were subjected to SDS PAGE and immunoblotting using an anti-GFP antibody. Size of the 250 kDa molecular weight marker is indicated. The asterisk denotes an unspecific band. C. Cell cycle distribution of the parental cell line and the TopBP1Δ284-301 cell line. The y-axis indicates the percentage of cells distributed in different cell cycle phases (G1, S, G2 and M) as indicated. The experiment was performed four times. For both the parental cell line and TopBP1Δ284-301, 1500 cells were analyzed by image cytometry for each experiment. Error bars represent the means ± 95% CI. D. *Upper panel*, western blot of cell extracts to detect CHK1-pS345. Extracts from the indicated cell lines were subjected to SDS PAGE and immunoblotting with an antibody against CHK1-Ser345-P. When indicated, cell cultures had been treated with 100 nM CPT for 30 min prior to harvest. *Lower panel*, ponceau staining to control for equal loading.

We used a cell line with a YFP-AID tag on all TopBP1 alleles (Pedersen et al., 2015) as parental background allowing for easy detection of TopBP1 in living cells. Using the outlined strategy, we obtained a number of clones with the desired deletion in the endogenous TopBP1 gene (Supplementary figure 1B). Using an antibody against the tagged TopBP1, we confirmed that the TopBP1 levels in the parental cell line and the TopBP1Δ284-301 cell line were similar (Fig. 2B) and that the deletion of the amino acids 284-301 does not alter the cell cycle distribution (Fig. 2C).

Moreover, by monitoring ATR substrate phosphorylation we found that TopBP1Δ284-301 is fully capable of activating ATR (Fig. 2D), which is not surprising given that the ATR activation domain is located between BRCT6 and BRCT7 and thus in another part of TopBP1. Yet this confirms that, the deletion of amino acids 284 to 301 of TopBP1 does not lead to an overall inactivation of TopBP1.

### The PLK1 docking site is important for the mitotic functions of TopBP1

Given the important functions of both TopBP1 and PLK1 in mitosis, we went on to investigate if the PLK1 docking site in TopBP1 affects the frequency of TopBP1 foci in mitosis. These foci reflect underreplicated DNA and other problematic DNA structures that need processing in mitosis to avoid transmission of DNA damage (Bagge et al., 2020; Pedersen et al., 2015). We thus performed live-cell fluorescence microscopy of untreated cells or cells treated with low dose aphidicolin (APH) for 16-20 hours, which induces mild replication stress and leads to elevated number of TopBP1 foci in mitosis (Pedersen et al., 2015). This analysis revealed that the TopBP1Δ284-301 forms more foci than the wildtype TopBP1 both with or without treatment with aphidicolin (APH) (Fig. 3A and B). Three other independently derived clones with TopBP1Δ284-301 also showed elevated levels of mitotic TopBP1 foci (Supplementary figure 1C). The reason for the increased number of mitotic TopBP1 foci can either be that the TopBP1Δ284-301 cell line enters mitosis with more underreplicated DNA and/or DNA damage or that it fails to reduce the number of TopBP1 foci during mitosis because of defective mitotic DNA repair. We also tested the effect of PLK1 inhibitor BI2536 on TopBP1 focus formation. PLK1 inhibition induces TopBP1 foci in the parental background consistent with the need for PLK1 activity to reduce TopBP1 foci (Fig. 3C). Importantly, PLK1 inhibition in the TopBP1Δ284-301 cell line does not lead to increase in TopBP1 foci (Fig. 3C) showing that this effect of PLK1 inhibition is fully dependent on the PLK1 docking site in TopBP1 that we have removed.

**Figure 3.**
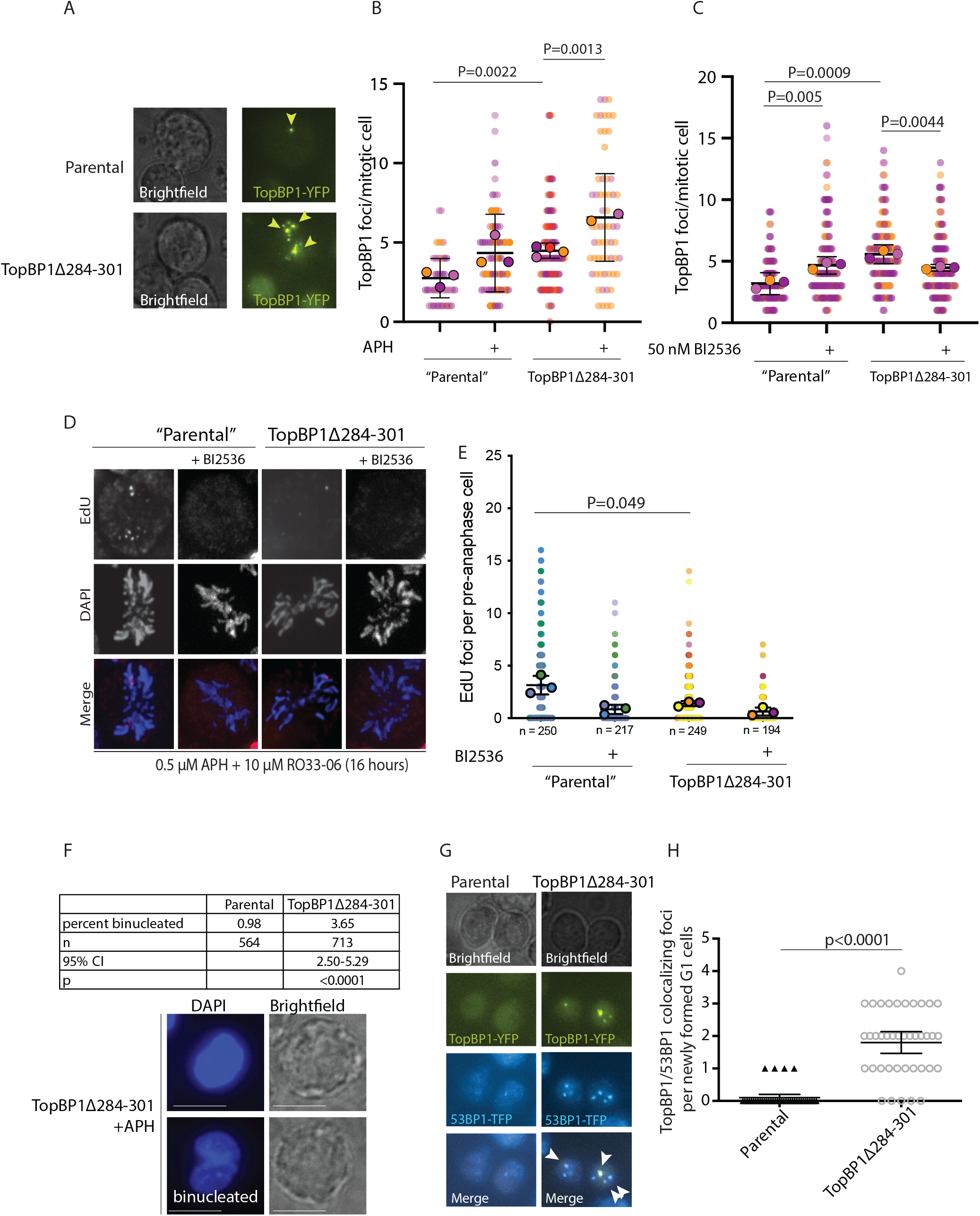
Characterizing the function of the TopBP1 PLK1 docking site in mitosis. A. Representative images of TopBP1^YFP-AID/YFP-AID/YFP-AID^ parental and TopBP1Δ284-301 cells. Brightfield (left) and fluorescence (right). TopBP1 foci in mitotic cells are indicated with yellow arrowheads. B. Superplot showing the quantification of TopBP1 foci per mitotic cell either untreated or treated with APH as indicated. Dots with a black border are the mean of each biologically independent experiment. Dots without a black border represent individual cells. Experiments are performed in triplicates represented by three different colors (the APH treated TopBP1Δ284-301 was done in duplicates) and statistical significance was calculated using the student’s t-test. Error bars represent the means ± 95% CI. *P*-values are indicated. C. Superplots showing the quantification of TopBP1 foci per mitotic cell untreated or treated with the PLK1 inhibitor BI2536. Dots with a black border are the mean of each biologically independent experiment. Dots without a black border represent individual cells. Experiments are performed in triplicates represented by three different colors. Error bars represent the means ± 95% CI. *P*-values are indicated. D. Representative images of EdU fluorescence and DAPI signal from pre-anaphase parental cells and TopBP1Δ284-301 cells. E. Superplots showing the quantification of EdU foci per pre-anaphase cell. Dots with a black border are the mean of each biologically independent experiment. Dots without a black border represent individual cells. Cells were treated with the PLK1 inhibitor BI2536 (50 nM) for 90 min where indicated. Experiments are performed in triplicates represented by three different colors. Error bars represent the means ± 95% CI. *P*-values are indicated. F. Percentage of binucleated cells is increased in the TopBP1Δ284-301 cell line. *Upper panel*, table showing percentage of binucleated cells of the parental and TopBP1Δ284-301 cell line. n refers to the number of cells analyzed. p, the statistical significance calculated using binomial test. *Lower panel*, images of DAPI fluorescence and brightfield of normal (upper) and binucleated (lower) cell. G. Representative images of mitotic TopBP1 foci colocalizing with 53BP1 in G1 phase. DT40 cells detected by time-lapse microscopy. The left part is parental cells and the right part is TopBP1Δ284-301 cells. Arrowheads mark TopBP1 foci that transitioned into 53BP1 NBs in G1. H. Quantification of the number of TopBP1/53BP1 colocalizing foci per early G1 cell. The statistical significance is calculated using student’s t-test. Error bars represent the means ± 95% CI.

MiDAS constitutes one mitotic DNA repair pathway that is facilitated by TopBP1 (Pedersen et al., 2015). We therefore investigated whether the mutant cell line was deficient in MiDAS. Our analyses show that EdU foci (representing MiDAS activity) were indeed much less abundant in the TopBP1Δ284-301 cell line compared to the parental cell line (Fig. 3D and E). Moreover, treatment of mitotic cells with a PLK1 inhibitor prior to EdU incubation severely reduces MiDAS in both the parental as well as the TopBP1Δ284-301 cell line (Fig. 3D and E) consistent with other studies (Minocherhomji et al., 2015; Wassing et al., 2021).

This indicates that the identified PLK1 docking site in TopBP1 is important for MiDAS and moreover suggests that the PLK1 TopBP1 interaction via the identified PLK1 docking site is facilitating the involvement of PLK1 in MiDAS.

We previously found that degron-induced depletion of TopBP1 leads to increase in the number of binucleated cells indicative of TopBP1 being involved in precise chromosome segregation (Pedersen et al., 2015). To investigate if the PLK1 docking site is important for this role of TopBP1 we compared the occurrence of binucleated cells in the two cell lines (Fig. 3F). Similar to TopBP1 depletion, the deletion of the TopBP1 PLK1 docking site results in an increased number of binucleated cells (Fig. 3F). Hence, we hypothesize that TopBP1 and PLK1 work together to achieve a faithful genome segregation and cell division.

Our results show that TopBP1Δ284-301 has a defect in MiDAS and chromosome segregation and we would expect that daughter cells of the TopBP1Δ284-301 cell line will have higher levels of DNA damage. To specifically address if this is the case, we set out to directly observe the extent of TopBP1 foci that develop into 53BP1 NBs in the M-G1 phase transition. Thus, we made endogenous 53BP1-TFP tagging in both cell lines using a previously described construct (Oestergaard et al., 2012) and the two resulting cell lines were subjected to time-lapse microscopy with an imaging frequency of 5 mins for a total of 50 mins. Although not all TopBP1 foci were trackable, we identified 40 cells containing trackable TopBP1 foci in each cell line. Quantification of TopBP1-53BP1 colocalizing foci in early G1 cells, shows that in contrast to the parental cell line most of the TopBP1Δ284-301 cells have TopBP1-53BP1 colocalizing foci in G1 (Figure 3G and H).

This strongly suggests that the mutant cell line fails to repair problematic DNA structures during mitosis and therefore pass on more DNA damage to daughter cells.

Taken together our analyses shows that the PLK1 docking site close to BRCT2 of TopBP1 is required to preserve genome integrity during cell division.

### The PLK1 docking site in TopBP1 is important cellular survival after PARP inhibition but not CPT treatment

All in all, our results show that the STP site in TopBP1 is important for the function of TopBP1 during mitosis. A previous study reported that PLK1-TopBP1 interaction facilitates RAD51 loading after irradiation of S-G2 cells (Moudry et al., 2016). Thus, we further investigated if TopBP1Δ284-301 impacts RAD51 loading during interphase. CPT treatment of replicating cells can induce one-ended DSBs that must be repaired by HR, thus RAD51 foci are expected to be induced by CPT treatment. We therefore performed RAD51 immunofluorescence experiments and the subsequent quantification revealed that the TopBP1Δ284-301 cell line has lower levels of CPT-induced RAD51 foci compared to the parental cell line (Fig. 4A and B).

**Figure 4.**
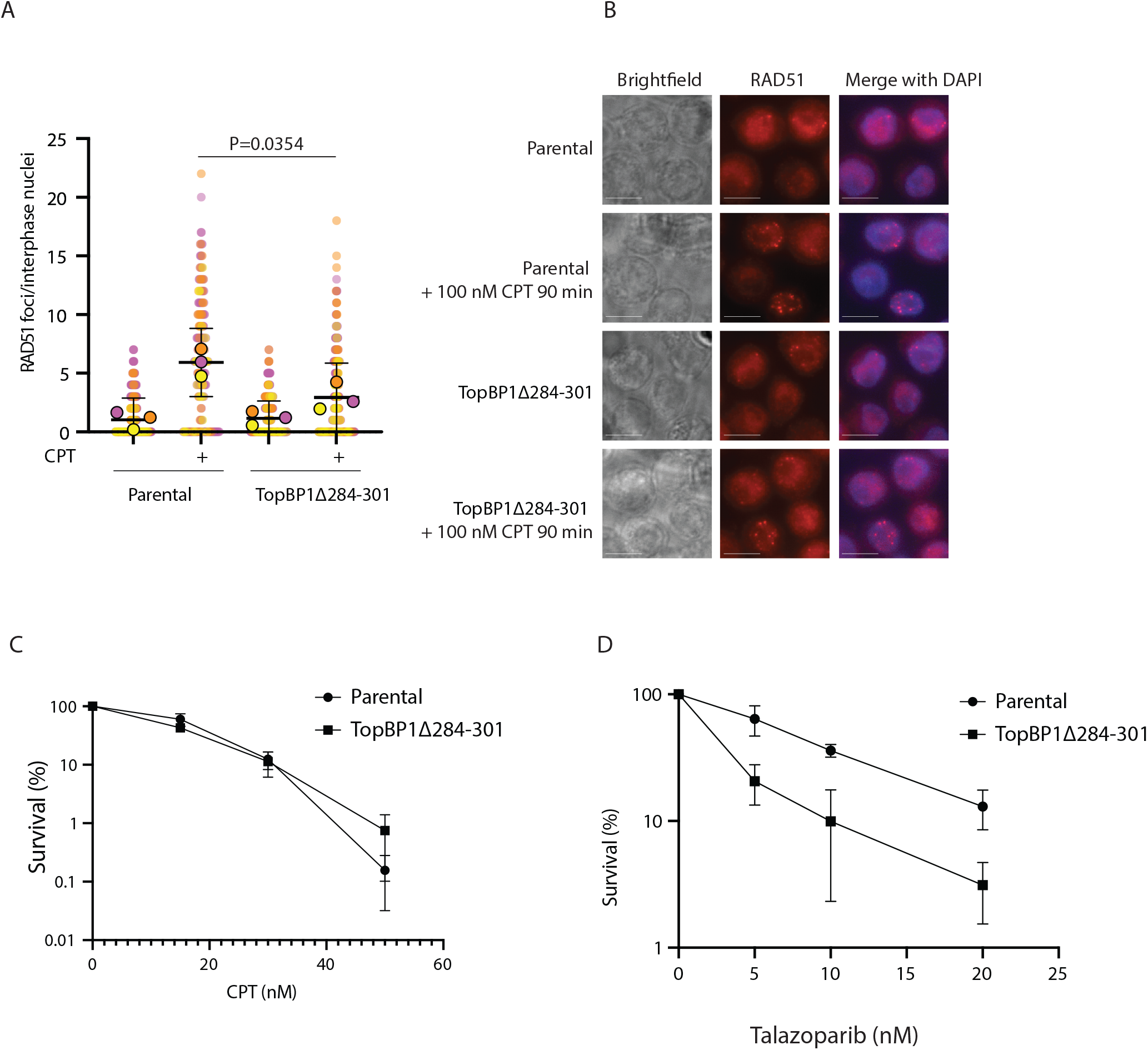
Investigating the function of the PLK1 docking in homologous recombination. A. Superplots showing quantification of the number of RAD51 foci per interphase nuclei. Dots with a black border are the mean of each biologically independent experiment. Dots without a black border represent individual cells. Experiments are performed in triplicates represented by three different colors. Statistical significance was calculated using student’s t-test. Error bars represent the means ± 95% CI. B. Representative images of RAD51 immunofluorescence and nuclei were stained by DAPI. Upper part is untreated or CPT treated parental cells and lower part is the TopBP1Δ284-301 cell lines with the same treatment. Scale bars represent 10 µm. C. Cellular sensitivity to CPT measured by colony survival assay of the two indicated cell lines in response to increasing CPT concentration. Experiments were performed in triplicates and statistical significance is calculated using student’s t-test. Error bars represent the means ± 95% CI. D. Cellular sensitivity to PARP inhibitor measured by colony survival assay of the two indicated cell lines in response to increasing CPT concentration. Experiments were performed in triplicates and statistical significance is calculated using student’s t-test. Error bars represent the means ± 95% CI.

The result shows that the deletion of the PLK1 docking site in TopBP1 leads to a decrease but not an abolishment of CPT-induced RAD51 focus formation. This indicates that homology-directed repair may be partially suppressed in the TopBP1Δ284-301 cell line. To address whether the suppression has a functionally significant effect on cellular survival after CPT treatment we employed colony survival assays. The result shows that the TopBP1Δ284-301 cell line is not hypersensitive to CPT (Fig. 4C). Given the dependence on homology-directed repair for survival of CPT induced DNA damage our results indicate that the TopBP1Δ284-301 cell line retains sufficient RAD51 loading activity to sustain homology-directed repair and cellular survival in response to CPT.

Since depletion of TopBP1 sensitizes cells to inhibitor of poly(ADP-ribose) polymerase (Moudry et al., 2016), we also tested by colony survival assays if the TopBP1Δ284-301 cell line is hypersensitive to the PARP inhibitor talazoparib. Interestingly, we found that deletion of the STP motif indeed sensitizes the cells to PARP inhibition (Fig. 4D) suggesting that the mitotic function of TopBP1 is important for survival after PARP inhibition but not CPT treatment.

## Discussion

We previously identified a function for the scaffold protein TopBP1 in mitosis. Specifically, we found that TopBP1 suppresses transmission of DNA damage to daughter cells (Pedersen et al., 2015). Importance of the mitotic function of TopBP1 was recently stressed due to the finding that the mitotic function of a CIP2A-TopBP1 complex is crucial for survival of BRCA negative cancer cells (Adam et al., 2021). This strongly suggests that the mitotic function of TopBP1 is a potential therapeutic target in BRCA deficient cancer cells, which is a strong incentive to gain better insight into regulation of the mitotic function of TopBP1.

There are several studies that propose a connection between TopBP1 and the mitotic kinase PLK1 (Moudry et al., 2016) (Balbo Pogliano et al., 2022) but the lack of specific mutations disrupting this interaction has hampered investigations into the significance of this. Here, we provide, to our knowledge, the first analysis of a TopBP1 mutant with a deletion of a conserved PLK1 docking site allowing us to address the function of this docking site in TopBP1 function during mitosis. Our results show that PLK1 binding to this site is required for TopBP1 functions during mitosis. Deletion of this site leads to increased transmission of DNA damage to daughter cells as well as increased frequency of binucleation.

Our finding that the PLK1 docking site on TopBP1 is important for the mitotic function of TopBP1 strongly suggest that this docking site brings PLK1 close to relevant substrates that must undergo phosphorylation to facilitate mitotic DNA processing. At this point it is unclear what those substrates are. However, serine 14 on RAD51 is an obvious candidate since previous studies have suggested that this residue is phosphorylated by PLK1 to facilitate MiDAS (Wassing et al., 2021). Also, we previously showed that TopBP1 is mediating this phosphorylation (Moudry et al., 2016). Moreover, recent work has shown that TopBP1 facilitates PLK1-mediated BLM phosphorylation (Balbo Pogliano et al., 2022). Thus, we find it likely that the identified PLK1 docking site orchestrates phosphorylation of several substrates important for DNA processing during mitosis. Of note, PLK1 has been suggested to activate SLX4, which is colocalizing with TopBP1 in mitosis (Bagge et al., 2020; Pedersen et al., 2015; Ragland et al., 2013). Finally, TopBP1 itself could also be substrate for PLK1 kinase activity. Determining the nature of the substrates as well as functional consequences of their phosphorylation will be subject of future studies.

It is worth noting that TopBP1 also contains two additional relatively conserved putative PLK1 docking site at residues 860-862 and at 1104-1106 in human TopBP1. These PLK1 docking sites might also be functional relevant and regulate other aspects of TopBP1 function, which will be important to establish in the future.

Finally, we find that the PLK1 docking site in TopBP1 is important for surviving PARP inhibitor treatment but not CPT treatment. Both of these drugs are thought to induce DNA damage that rely in homologous recombination for repair. A hypothetical explanation for this finding is that lesions induced by PARP inhibition may be more likely to escape checkpoint detection and persist into mitosis than CPT induced DNA damage and that the PLK1 docking site that we have described is critically important for DNA repair in mitosis but not for homologous recombination in interphase cells.

All in all, this study sheds light on the regulation of TopBP1 during mitosis, which may be important for developing new strategies to target this pathway in BRCA deficient cancers.

## Materials and methods

### Protein expression and purification

His-tagged PLK1 was expressed in the *E. coli* strain BL21 Rosetta2 (DE3) R3 T1 at 18°C for 20 h using 0.5 mM IPTG. The bacterial pellets were resuspended in ice-cold buffer L (50 mM NaP, 300 mM NaCl, 10% Glycerol, 0.5 mM TCEP, pH 7.5) containing complete EDTA-free Protease Inhibitor Cocktail tablets and lysed with an EmulsiFlex-C3 High Pressure Homogenizer. The lysate was centrifuged at 18,500 × *g* for 30 min and the supernatant filtered through a 0.22 µm PES filter and loaded onto a 1 mL Ni column (GE healthcare) in buffer L with 10 mM imidazole, washed and eluted. The eluate was loaded on a Superdex 200 PG 16/60 equilibrated with SEC buffer (50 mM NaP, 150 mM NaCl, 0.5 mM TCEP, 10% Glycerol, pH 7.50) and fractions were analyzed by SDS-PAGE and verified by mass spectrometry.

### Isothermal Titration Calorimetry (ITC)

Peptides were purchased from Peptide 2.0 Inc (Chantilly. VA, USA). The purity obtained in the synthesis was 95 – 98% as determined by high performance liquid chromatography (HPLC) and subsequent analysis by mass spectrometry. Prior to ITC experiments both the protein and the peptides were extensively dialyzed against 50 mM sodium phosphate, 150 mM NaCl, 0.5 mM TCEP, pH 7.5. All ITC experiments were performed on an Auto-iTC200 instrument (Microcal, Malvern Instruments Ltd.) at 25°C. Both peptide and Plk1 concentrations were determined using a spectrometer by measuring the absorbance at 280 nm and applying values for the extinction coefficients computed from the corresponding sequences by the ProtParam program (http://web.expasy.org/protparam/). The peptides at approximately 450 μM or 120 μM (for submicromolar affinities) concentration were loaded into the syringe and titrated into the calorimetric cell containing the Plk1, respectively, at ~ 35 μM or ~ 10 μM. For the ggTopBP1 double phosphorylated peptide, ggTopBP1^287-300^ pT291, pT296, a concentration of 60 μM in the syringe was used. The reference cell was filled with distilled water. In all assays, the titration sequence consisted of a single 0.4 μl injection followed by 19 injections, 2 μl each, with 150 s spacing between injections to ensure that the thermal power returns to the baseline before the next injection. The stirring speed was 750 rpm. Control experiments with the peptides injected in the sample cell filled with buffer were carried out under the same experimental conditions. These control experiments showed heats of dilution negligible in all cases. The heats per injection normalized per mole of injectant *versus* the molar ratio [peptide]/[Plk1] were fitted to a single-site model. Data were analysed with MicroCal PEAQ-ITC (version 1.1.0.1262) analysis software (Malvern Instruments Ltd.).

### Cell culture

DT40 cells were cultured in RPMI 1640 medium GlutaMAX (ThermoFisher Scientific) supplemented with 2% chicken serum (Sigma-Aldrich), 8% fetal bovine serum (ThermoFisher Scientific), 50 μM β-mercaptoethanol, 50 U/ml penicillin, and 50 μg/ml streptomycin at 37°C with 5% CO_2_. HeLa cells were maintained in DMEM (ThermoFisher Scientific) with 9% FBS (ThermoFisher Scientific) and 50 U/ml penicillin, and 50 μg/ml streptomycin and grown at 37°C with 5% CO_2_.

### Genetic targeting in DT40 cells

Plasmids and primers used in this study are listed in Supplemental Table S1 and S2. Target sites for Cas9 were identified using the Benchling online tool and targeting construct were constructed as described by Ran et al. (Ran et al., 2013) using pX458 as a backbone. This plasmid allows for expression of guide RNA, Cas9 nuclease and GFP. The sequences for DNA oligoes with target sites are given in table 1. The correct insertion of target sites was validated by PCR followed by sequencing (Eurofins). For a two target site deletion strategy, 7.5 µg of each targeting vector were transfected into DT40 using Amaxa transfection as previously described (Franklin and Sale, 2006). At day one after transfection, GFP positive cells were single-cell sorted into 96 well plates. Clones were scaled up and screened with primers flanking the target site. PCR products were analysed by gel electrophoresis and Sanger sequencing. Targeted transfection to obtain endogenous tagging of 53BP1 in DT40 cells was carried out as previously described (Oestergaard et al., 2012; Pedersen et al., 2015).

### Colony survival assay

Methyl cellulose medium was prepared as previously described (Simpson and Sale, 2006). For colony survival assays with BI2536 (Selleck-chemicals), talazoparib (Cat#S7048, SMS-Gruppen Denmark) or CPT (Cat#C9911, Merck), the methylcellulose medium with drug was shaken for more than 1 hour at 4°C before transfer into 6 well plates with 5 ml medium each well.

### Immunostaining

For interphase Hela cells the poly-lysine covered coverslip was added to wells in 24-well plate and 4×10^5^ cells/ml in 1 ml of DMEM medium was added. On the next day, medium was removed and the cells were washed with PBS before proceeding with immunostaining.

To stain mitotic HeLa, cells were treated 0.1 μg/ml colcemid (Life Technologies) for 6 hours to arrest cells in mitosis. Then the mitotic cells were shaken off and harvested at 1000 rpm for 5 min.

For staining of DT40, exponentially growing cells were harvested and resuspended in PBS. 100 μl with 2.5×10^6^ cells were added onto one poly-lysine coated coverslips and left for 10 min before proceeding directly to the immunostaining.

Immunostaining of all cell types were performed as described here. 400 μl 3% paraformaldehyde was carefully added to each coverslip and the slides were incubate at room temperature for 18 min. Then, the supernatant was removed and cells were washed three times with 400 μl PBS. Subsequently, 400 μl freshly prepared 0.1% Triton-X-100 in PBS-T was used to permeabilize cells for 18 min after which all liquid was removed and the coverslips were washed three times with 400 μl PBS. Then 400 μl glycine (25 mM) was added for 7 min before washing twice with 400 μl PBS. 400 μl 3% BSA in PBS-T (blocking solution) was added onto each coverslip and incubated at 4°C overnight. Coverslips were incubated with 200 μl primary antibody in blocking solution at room temperature. The primary antibody was removed and coverslips were washed four times 10 min with 400 μl PBS-T (PBS with 0.1% Tween-20). Afterwards, coverslips were incubated with 200 μl secondary antibody-solution, and incubated at 37°C for 45 min, and then the coverslips were incubated with secondary antibody at room temperature for 1 hour protected from light. Finally, coverslips were washed 4 times 10 min with 400 μl PBS-T and washed with double distilled water immediately before mounting with DAPI mounting medium (4% *n*-propyl gallate, 80% glycerol, 1 μg/ml DAPI). Primary antibodies were used in the following dilutions in 0.1% PBS-T, 3% BSA: rabbit anti-Rad51, Bio Academia, Cat#70-001 1:1000; mouse anti-PLK1, Santa Cruz, Cat#sc-17783, 1:200; rabbit anti-TopBP1, Cat#A300-111A, 1:200; Bethyl Laboratories, Inc., 1:1000. The following secondary antibodies were used in 1:1000 dilutions in 0.1% PBS-T, 3% BSA: Alexa594 anti-mouse, Life Technologies, Cat#A-11005; Alexa594 anti-rabbit, Life Technologies, Cat#A-11037; Alexa 488 anti-mouse, Life Technologies, Cat#A-21121; STAR-RED anti-rabbit, Abberior, #STRED-1002.

### Binucleation assay

On day 1, cell cultures were or were not treated with 0.4 μM APH at 37°C for 16-20 hours. On day 2, untreated and treated cells were harvested and mounted with DAPI mounting medium, as explained for immunostaining except that the primary and secondary antibody incubation was omitted.

### Microscopy

The DeltaVision Elite Microscope from GE Healthcare Life Sciences with a 100X lens and the SoftWorx 7.0.0 software was used to image DT40 and Hela cells. For the sectioning, optical section spacing was 0.5 μm, and the number of optical sections was 6. Subsequently, Volocity software from PerkinElmer was used to analyze images. Finally, all graphs and statistical analyses were performed in GraphPad Prism 8 software and superplots prepared according to the article (Lord et al., 2020)

### Live-cell microscopy of TopBP1 foci in DT40 cells

For live cell microscopy of DT40 cell lines, exponential growing cells in RPMI medium were used. Cells were untreated or treated with drugs as specified here: 0.4 µm APH for 16-20 hours or 50 nm BI2536 for 1.5 hours. Cells were imaged in an Ibidi µ-Slide Angiogenesis. The time-lapse microscopy was performed with an imaging frequency of 5 min for 50 min total.

### Detection of mitotic DNA synthesis (MiDAS)

1.6·10^6^ cells from an exponentially growing culture were pelleted at 1000 rpm for 5 minutes. Hereafter, cell pellets were resuspended in 2 ml media containing RO-3306 (10 µM) and Aphidicolin (0.4 µM) to a total cell density of 0.8·10^6^ cells/ml. The cell suspension was placed in a 6 well plate for 16 hours in incubator at 37°C to synchronize cells at the G2/M border. The next day, the cells were pelleted by centrifugation at 1000 rpm for 5 minutes. The cells were resuspended in 10 ml fresh pre-warmed media, to wash away RO-3306, and pelleted by centrifugation at 1000 rpm for 1 minute. The pelleted cells were then resuspended in media containing EdU (20 µM). Alternatively, cells were resuspended in media containing EdU (20 µM) and PLK1-inhibitor (BI-2536, 100 nM). The cells were pulsed with EdU or EdU+PLK1i for 35 minutes in the incubator. Following the EdU-pulse, 200 µl of cell suspension were spun onto a glass slide using the Cytospin centrifuge (Thermo) at 1000 rpm for 5 minutes.

The cells were fixed onto the glass slide for 10 minutes using 4% formaldehyde. Hereafter, samples were placed in a Coplin jar and permeabilized for 10 minutes in a KCM buffer (120 mM KCl, 20 mM NaCl, 10 mM Tris-HCl pH 8.0, 0.5 mM EDTA, 0.1% Triton X-100). After permeabilization, slides were washed twice in PBS-T for 5 minutes. Next, slides were washed once in 3% BSA (PBS-T) for 10 minutes. Hereafter, EdU was labelled using the 647 azide plus kit (Click-&-Go Plus EdU 647, Cat#1353, Click Chemistry Tools) according to the manufacturers protocol. Slides were then washed once in PBS-T for 5 minutes and once in PBS for 5 minutes. 10 µl of Prolong Diamond containing DAPI (ThermoFisher, Cat#P36962) were added on top of the cells and covered with a coverslip and sealed with transparent nail-polish. Slides were kept at 4°C until imaging.

### Western blot

Western blots were performed as previously described (Shao et al., 2020). For analysis of TopBP1-YFP-AID mouse anti-GFP monoclonal (1:500, Cat#11814460001, Roche) and anti-mouse (IRDye® 800CW Goat anti-Mouse IgG, Cat#926-32211, LI-COR) were used as primary and secondary antibodies, respectively. For analysis of CHK1-Ser345P, rabbit anti-phospho Ser345-CHK1 (1:1000, Cat#2348, Cell Signaling), and anti-rabbit IgG conjugated to HRP (1:2000, Cat#P0217, Dako) were used as primary and secondary antibodies, respectively. Development of the western blot was performed using ECL (Cat#RPN2236, Amersham) according to manufacturer’s instruction. Alternatively, for the IR dye the blot was visualized using a LI-COR instrument.

### Cell cycle analysis

For measuring the percentage of H3 phospho-Ser10-positive DT40 cells, cells were harvested and washed with ice-cold PBS, fixed with 70% EtOH at −20°C for overnight, followed by two PBS washes, and incubation with pS10H3 antibody solution (1:40, Cat#06-570, Millipore) at room temperature for 2 h. This was followed by incubation with secondary antibody (Alexa 568 goat anti-rabbit, Cat#A11011, Life Technologies) for 30 min at room temperature in the dark. Finally, cells were resuspended in 300 μl 0.1% Triton X-100 in PBS with 1 μg/ml DAPI and left for 5 min at 37°C before transfer of 40 μl of stained cell culture into an Xcyto 2-chamber slide. Images were processed and quantitative measurement of fluorescence intensities was performed using Xcyto 10 (ChemoMetec).

## Acknowledgements

This work was supported by the Villum Foundation to V.H.O. and M.L., the Danish National Research Foundation (DNRF115) to M.L., and the Novo Nordisk Foundation to J.B., M.L. and V.H.O. Work at the Novo Nordisk Foundation Center for Protein Research is supported by grant NNF14CC0001. We thank the protein production and characterization unit at NNF CPR for producing Plk1 polobox domain. We thank Blanca Lopez Mendez from the protein production and characterization unit at NNF CPR for performing the ITC measurements. We thank Arminja Kettenbach for comments on PLK1 interactomes and Norman Davey for prediction of putative PLK1 docking sites in PLK1 interactors.

