## supplementary for "A conserved PLK1 docking site in TopBP1 maintains genome integrity during mitosis"

**Table S1.** Cell lines

| Cell type | Cell line genotype | Source |
|---|---|---|
| DT40 | TopBP1-YFP-AID | (Pedersen et al., 2015) |
| DT40 | TopBP1Δ284-301 cl.1, cl.25, cl.40, cl.41 | This study |
| DT40 | TopBP1-YFP-AID, 53BP1-TFP | This study |
| DT40 | TopBP1Δ284-301 TopBP1-YFP-AID, 53BP1-TFP | This study |
| HeLa | “WT” | |

**Table S2.** DNA oligos

| Name | Sequence | Use |
|---|---|---|
| JL9 | CACCGACAATGTATAAAATAGAGAG | Target site for CRISPR-Cas9 mediated deletion of TopBP1 PLK1 docking site |
| JL10 | AAACCTCTCTATTTTATACATTGTC | Target site for CRISPR-Cas9 mediated deletion of TopBP1 PLK1 docking site |
| JL11 | CACCGTACTGTCAAGCTTGCTGGCG | Target site for CRISPR-Cas9 mediated deletion of TopBP1 PLK1 docking site |
| JL12 | AAACCGCCAGCAAGCTTGACAGTAC | Target site for CRISPR-Cas9 mediated deletion of TopBP1 PLK1 docking site |
| JL15 | AGGGGGCTTCTTCGTGCTTT | Screening of TopBP1 PLK1 docking site deletion |
| JL16 | CGTAAGCAAAGCGGAAACCA | Screening of TopBP1 PLK1 docking site deletion |
| RTP35 | AGCACTAATCCCTGAGGAATGCCAC | Screening of endogenous tagging of 53BP1-TFP in DT40 cell line |

| | | |
|---|---|---|
| RTP34 | GTCTTGTAGTTGCCGTCGTCCTTGA | Screening of endogenous tagging of 53BP1-TFP in DT40 cell line |

**Table S3.** Plasmids

| Name | Use |
|---|---|
| px458 (pSpCas9(BB)) (Ran et al., 2013) | Used for CRISPR-Cas9 mediated KO of TopBP1 PLK1 docking |
| pCR2.1gg53BP1-TFP (Oestergaard et al., 2012) | transfected the 53BP1-TFP knock-in constructs into DT40 YFP-AD/YFP-AID/YFP-AID cells |

#### Figure legend

##### Supplementary figure 1

A. Isothermal titration calorimetry binding curves for the interaction between the indicated peptides from human or chicken TopBP1 and PLK1.

B. Traces from Sanger sequencing of successfully targeted clone with a deletion of the genomic region encoding TopBP1 amino acid 284-301 on all alleles. Position of the deleted region is indicated with a dashed line and the 54 bases including conserved STP encoding part is shown above in blue.

C. Quantification of TopBP1 foci per mitotic cell of the parental cell line and four independently derived clones with TopBP1Δ284-301. The red bar indicates the mean.

### Supplementary figure 1

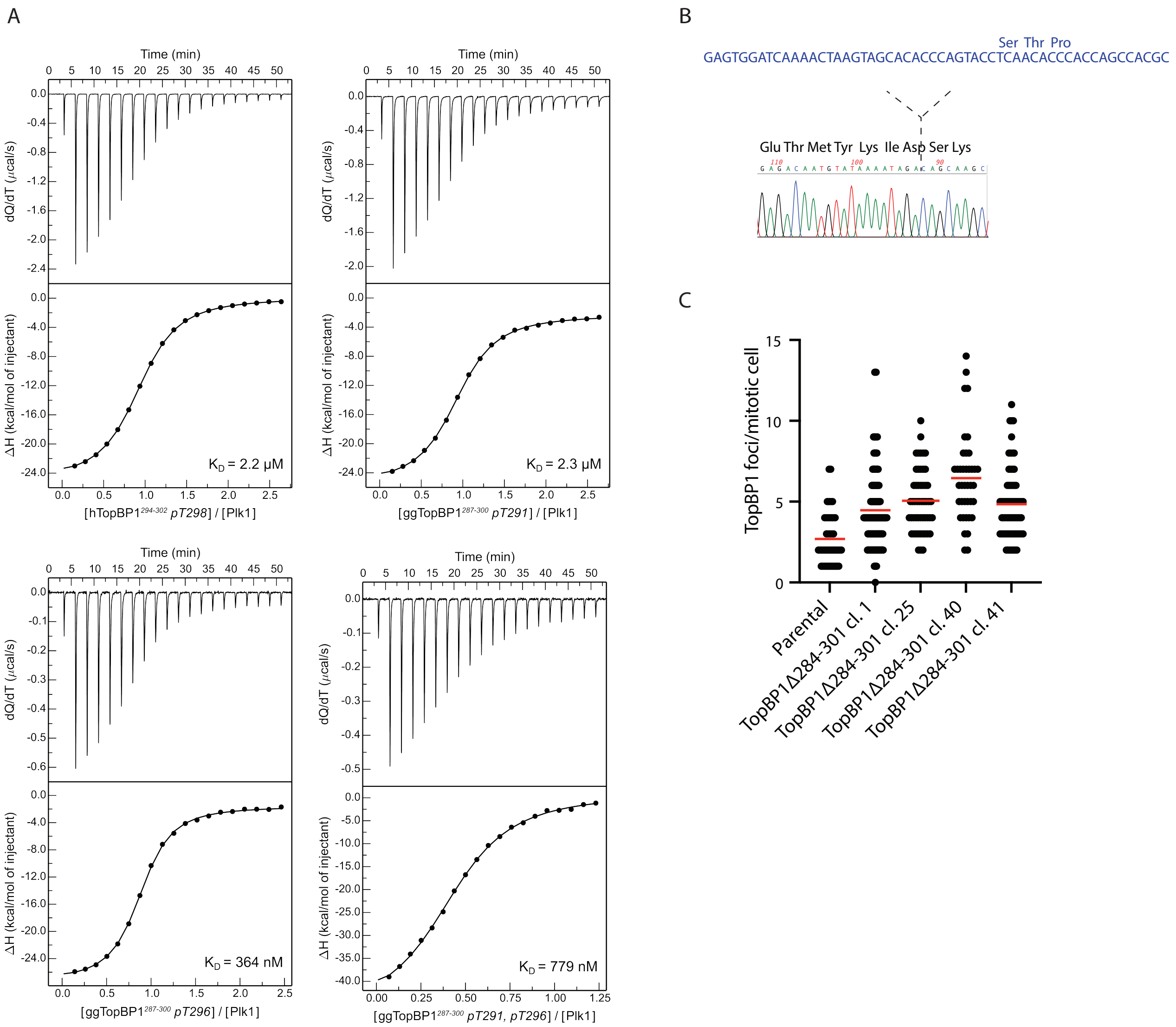
